# Individual variation in tolerance of human activity by urban dark-eyed juncos (*Junco hyemalis*)

**DOI:** 10.1101/2020.12.11.419937

**Authors:** Hayley M. Stansell, Daniel T. Blumstein, Pamela J. Yeh, Peter Nonacs

## Abstract

An important goal of urban ecology is determining what differentiates urban-tolerant populations of birds from their non-urban ancestors and urban-intolerant species. One key to urban success may be reacting appropriately to human activity, and the degree to which birds view humans as threats can be quantified by their escape behavior. Understanding individual-level plasticity, however, requires the tracking of known individuals. We compared flight-initiation distances (FID) and distances fled (DF) from approaches by a human across urban and non-urban populations of individually-marked Dark-eyed Juncos (*Junco hyemalis*) in southern California. The urban population is more tolerant to people as evidenced by attenuated FIDs and DFs relative to non-urban birds. Although individual urban birds either habituated or sensitized to repeated approaches, there was no significant pattern at the population level. Overall, the behavioral patterns exhibited by urban juncos are more supportive of *in situ* evolution than either being a biased sample from an ancestral non-urban population or intrinsic behavioral plasticity that produces a uniform adjustment to urban life.

## INTRODUCTION

Urbanization poses a rapidly-growing threat to wildlife worldwide, including native birds as it affects food availability, predation pressure, and habitat quality (Chace and Walsh 2006). Furthermore, urban settings contain stressful and generally detrimental stimuli that include more noise, pollution, and human activity. Nevertheless, some species prosper in urban environments while others suffer or are pushed out entirely (Chace and Walsh 2006, Schlesinger et al. 2008). Understanding how species adapt and change to survive in cities can inform conservation and urban planning decisions to support the maintenance of native biodiversity in proximity to human activity (Fernández-Juricic et al. 2001, Chace and Walsh 2006, Aronson et al. 2014).

Direct disturbance by human activity may be one of the primary stressors faced by urban birds (Partecke et al. 2006, Strasser and Heath 2013). One way to understand the cost of human disturbance on wildlife is based on the premise that escape behavior shows how wildlife perceive humans as a risk to avoid (Frid and Dill 2002, Blumstein 2013). Escape decisions can vary depending on the economics of fleeing, lost opportunity cost, and perceived risk of predation (Ydenberg and Dill 1986, Cooper and Blumstein 2015). All of these factors could differ for urban and non-urban birds, leading to consistent differences observed in escape behavior (Møller 2008, Evans et al. 2010, Mikula 2014, Samia et al. 2015, Sprau and Dingemanse 2017).

Recently-established urban populations of Dark-eyed Juncos (*Junco hyemalis*) have undergone rapid changes in physiology, morphology, and behavior. These have occurred over the course of a few decades (Yeh 2004; Yeh and Price 2004; Newman et al. 2006; Atwell et al. 2012; Atwell et al. 2014). One change that may allow juncos to colonize and then thrive in cities is in increasing tolerance to human activity. Such tolerant urban populations may arise in several ways (Sol et al. 2013). First, urban populations may result from differential habitat selection by intrinsically more human-tolerant subsets of individuals from non-urban ancestor populations (Carrete and Tella 2011). Second, urban birds may have evolved to be more tolerant of disturbance (Møller 2008, Carrete et al. 2016). Third, urban species are behaviorally more plastic and thus better at appropriately habituating to human activity (Fernández-Juricic et al. 2001, Rodriguez-Prieto et al. 2009, Møller 2010, Blumstein 2016, Vincze et al. 2016). Here, our goal is to differentiate between inter-individual variation and within-individual plasticity in urban birds. We gathered repeated samples in juncos of flight-initiation distance (FID) and distance fled (DF) over both short time intervals (within-day) and longer intervals (across-days). For a baseline comparison of urban to non-urban juncos, we also measured FID and DF values from a nearby non-urban population.

## METHODS

### Study Sites and banding birds

The urban site was the 170ha campus of the University of California Los Angeles (hereafter: UCLA, 34.0695°N, 118.4452°W). Abundant human activity occurs throughout the year, although pedestrian activity fluctuates daily, weekly, and monthly. The campus contains lawn and a mix of largely non-native plant species. The non-urban site was the James Reserve (in the San Jacinto Mountains, 33.8083°N, 116.7778°W). Compared to UCLA, the 20 ha of the reserve and adjacent areas have substantially lower pedestrian activity. The habitat includes montane riparian forest and mixed conifer and hardwood forest with open understory consisting of gravel roads, parking areas and grassy meadows.

Mist-netted birds were individually marked at each site with USGS aluminum bands and a unique set of color bands. After banding, birds were given at least a week to recover before any behavioral data were taken. Mated pairs were never simultaneously tested with at least a week between taking data on the first and second bird.

### Sampling

All encounters were recorded by the same individual (HMS) following the protocol in commonly used to study FID (Cooper and Blumstein 2015). An encounter began when a bird was observed foraging or stationary, and not alarm calling or otherwise visibly agitated. Because juncos are territorial, most encounters with a given bird occurred near where it was initially captured and banded. Birds were always approached in a straight line at a practiced pace (approximating 0.5m/sec) when on the ground and exposed from vegetation, with no obstacles or other juncos between the observer and the focal bird. This ensured consistent, readily-detectable approaches to each individual (Frid and Dill 2002, Tätte et al. 2018). A colored marker was dropped at the location where the experimental approach began (the starting distance, SD), a second at the observer location when the focal subject fled (the FID), and a third at the location from where the focal subject fled, later converting paces to meters (0.825m/step). The distance fled (DF) was recorded by visually estimating the horizontal and vertical distance travelled in meters, then converting to a Euclidian distance. In some cases, it was not possible to collect data on DF because the bird left the immediate area. Such occurrences were arbitrarily recorded as a distance fled of 50 m.

We also recorded the time of day, presence/absence of conspecifics within a 5 m radius of the focal bird, distance to nearest cover, and pedestrian density. Pedestrian density (only at UCLA – there were almost never pedestrians at the James) was recorded categorically as low (defined as < 5 people per minute crossing a 10 m sample transect in the immediate vicinity of the approach) or moderate to high (≥ 5 people/min).

At UCLA, we collected repeated measures for 22 series on both a short time scale (4 attempted approaches to the same bird on the same day) and on a longer time scale (over a consecutive 4-day period). Not every individual had the complete sequence of 16 approaches (6 had one missing value in terms of a missed approach on one day, 4 had two missing values, and 1 had four missing values because of a missed day). Birds were approached during the 2017 breeding season while rearing chicks (February to July at UCLA; June to July at the James). In total, we collected 404 approaches across 31 individuals at UCLA, and 104 approaches across 22 individuals at the James Reserve. At both sites, the majority of data were collected between 08:00 and 13:00 h.

Statistical tests on the multiple approaches towards marked birds in UCLA followed standardized analytical methodology (Pezner et al. 2017, Dehaudt et al. 2019, Andrade and Blumstein, 2020). All analyses used R 3.4.2 (R Core Team 2017), on code modeled after Pezner et al. (2017), which had an identical experimental design: 4 trials per day, repeated across 4 consecutive days. We used linear (FID) and logistic (DF, with distances ≤2 m as near and >2 m as far) mixed-effects models for individual responses to repeated approaches on the UCLA campus.

We fitted models using the R package “lme4” v1.1-14 (Bates et al. 2015) (supporting package “car” v2.1-6: Fox and Weisberg 2011). A null model used individual bird as a random intercept, then iteratively incorporated fixed effects (contextual variables), with stepwise selection to find the combination of fixed effects with the lowest AIC value. Each predictor variable was added to the model and then selectively removed depending on their effect on AIC relative to the null model. In cases where a fixed effect resulted in only a non-significant decrease in AIC, likelihood ratio tests used “lmerTest” v2.0-33 (Kuznetsova et al. 2016) to measure significance. Fixed effects without significant model improvement were discarded.

After selecting a model via this process, a likelihood ratio test evaluated whether either the inclusion of trial iteration as a fixed effect or as a random slope significantly improved the explanatory power over the model containing contextual predictor variables and the random intercept. Best mixed-models were compared against their fixed effects-only counterparts via likelihood ratio test using “RLRsim” v3.1-3 (Scheipl et al. 2008), supporting packages “MASS” v7.3-47 and “arm” v1.9-3 (Gelman and Su 2016) to determine whether individual differences among birds explained a significant portion of behavioral variation. Where individual was a significant random effect in models, adjusted repeatability was calculated using code provided by Jean-Nicolas Audet, modified from “rptR” v0.9.21 (Stoffel et al. 2017). Analyses were based on 22 series of approaches, with 334 individual approaches.

## RESULTS

### Individual Variation in the Urban Population

A variety of factors when directly examined do not significantly explain variation in FID. These include distance from cover (linear regression: *F*_1,369_ = 2.573, *R*^*2*^ = 0.004, *p* = 0.110), sex (male mean FID = 3.23 m ± 1.06 (sd), n = 17; female mean FID = 3.72 ± 1.81, n = 6; *t* = 0.809, *p* = 0.428), and presence/absence of conspecifics with 5 m (within bird matched-pair *t* = 0.349, *p* = 0.731, n = 19). Interestingly, birds fled at a greater distance when pedestrian density was low than when it was moderate or high, but the effect did not quite reach statistical significance (low: mean FID = 3.52 ± 1.38; moderate/high mean FID = 3.13 ± 1.48; within bird matched-pair *t* = 2.013, *p* = 0.058, n = 21).

For within-day mixed effects models of FID, starting distance was also the only fixed effect retained in mixed models across days (Supp. Table 1). Increasing starting distance was associated with an increased FID. Increasing FID was associated with an increased DF on both time scales. Distance to cover, retained as a fixed effect, was not significant with a change in DF over short time scales (Supp. Table 2).

Trial number explains neither significant variation in FID (Supp. Tables 3 and 4), nor DF (Supp. Tables 5 and 6) within or across days. Trial number as either a fixed effect or a random slope failed to significantly improve model fit compared to a random-intercept only model. Therefore, repetition in flushes does not significantly affect FID or DF as would be predicted by either all birds habituating or sensitizing.

Likelihood ratio tests comparing mixed models against linear models indicated a significant individual bird effect (*p* < 0.01 for both time scales) on FID. Similarly, adjusted repeatability tests suggest a large proportion of the variation in FID, but not in DF, is explained by individual bird after controlling for fixed effects (*R*^*2*^ = 0.46 within-day, *R*^*2*^ = 0.43 across-days, Supp. Tables 1, 5 & 6).

FID slopes within and across days (Fig. 1) are significantly positively correlated across series, but DF slopes are not (FID: *F*_1,20_ = 4.886, *R*^*2*^ = 0.156, *p* = 0.039; DF: *F*_1,20_ = 0.690, *R*^*2*^ < 0.001, *p* = 0.416). Thus, birds which increased FID with each approach within a day also tended to increase FID with each subsequent day they were approached. Individual FID slopes did not significantly predict DF slopes (slopes within-day: *F*_1,20_ = 0.404, *R*^*2*^ < 0.001, *p* = 0.532; slopes across-days: *F*_1,20_ = 1.252, *R*^*2*^ = 0.012, *p* = 0.276). Some birds significantly decreased FID over repeated flushes (consistent with habituation), while others increased FID (consistent with sensitization; Fig. 2).

**FIGURE 1:**
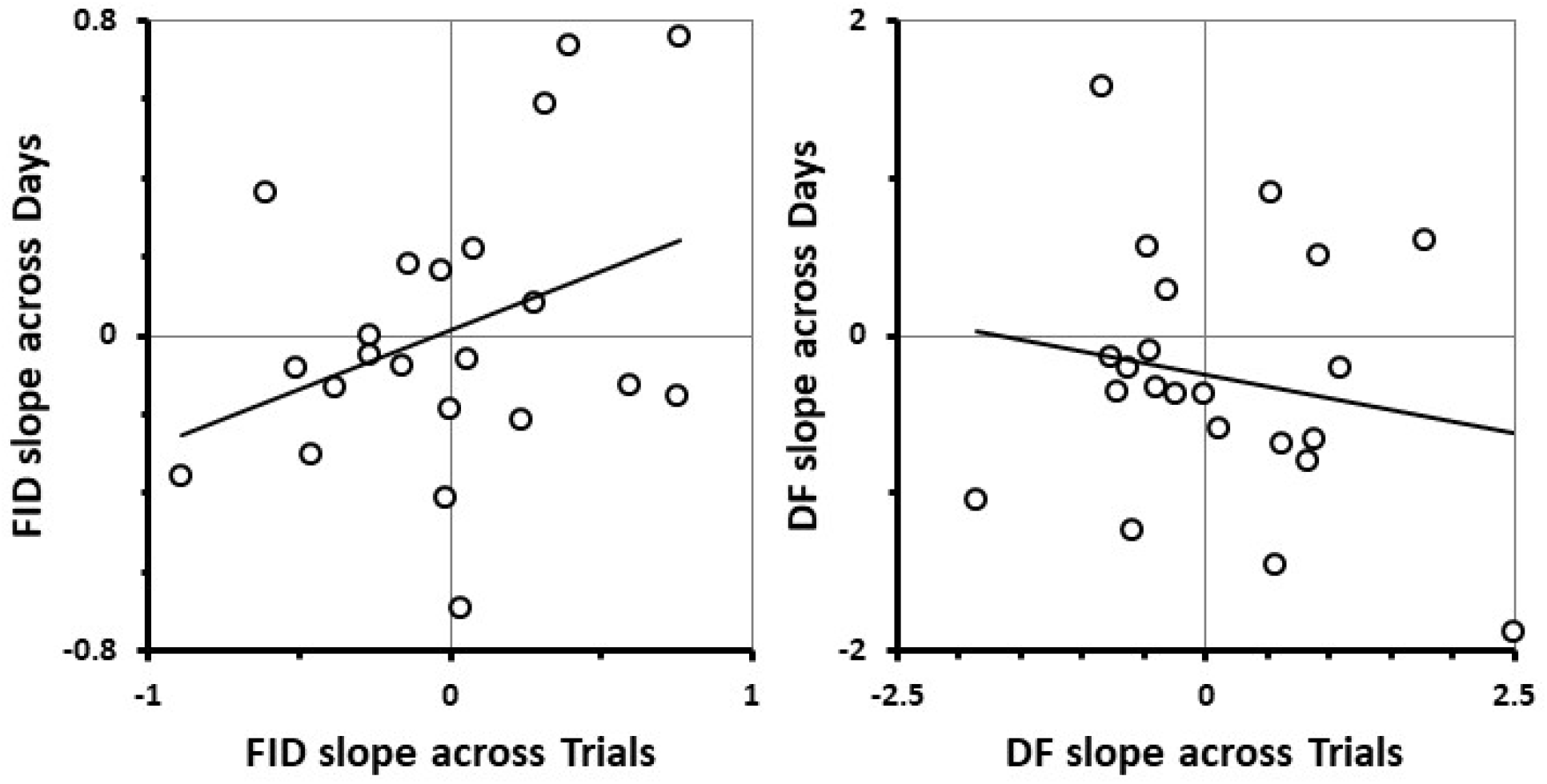
Regressions of FID and DF slopes. Each data point (n = 22) represents one series of approaches. Slopes for Trials are the average of the four individual slopes for each consecutive approach (i.e., the four slopes of the changes in FID or DF from day 1 to day 4 for approaches #1, 2, 3, or 4). Slopes for Days are the average of the four individual slopes for each consecutive day (calculated as the change in FID or DF from the first to fourth approach of that day).

**FIGURE 2:**
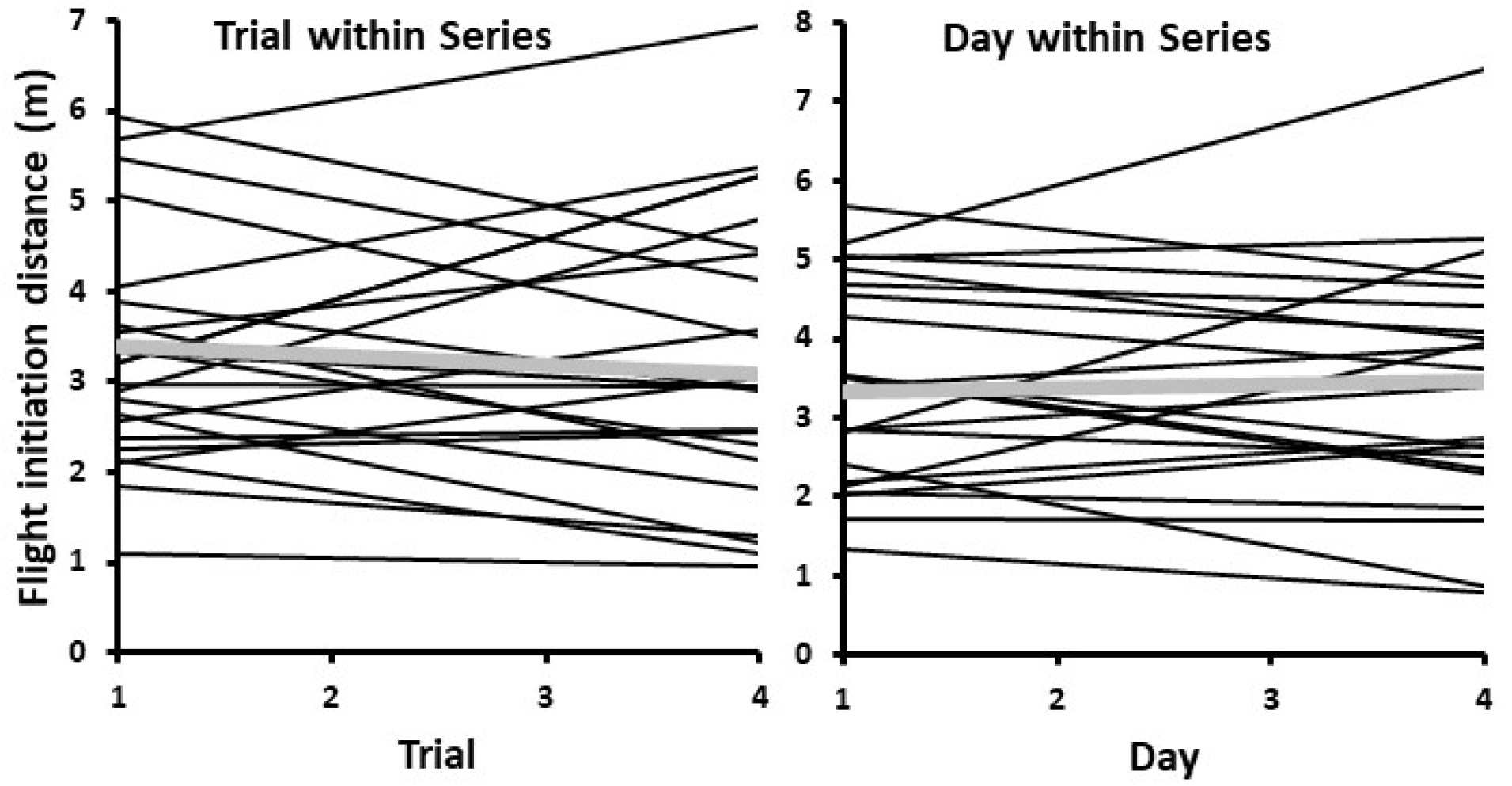
Individual urban junco FIDs by trial and day. Positive slopes indicate sensitization, negative ones indicate habituation. Thin black lines indicate individual series (n = 22), while the thick grey lines show the mean across all the series.

### Comparison of Urban and Non-Urban Populations

In comparing their first flushes, urban juncos had both significantly shorter and less variation FIDs than non-urban birds (Fig. 3: urban = 3.49 ± 1.47 m (sd), n = 31; non-urban = 9.67 ± 4.29 m, n = 22; *t* = 7.46, *p* < 0.001). Urban birds were encountered closer to cover than non-urban birds (1.36 ± 1.51m versus 4.23 ± 2.12m, *t* = 5.64, *p* < 0.001). This likely accounts for the shorter mean distances fled by urban juncos relative to non-urban (Fig. 3: 4.94 ± 4.63 m versus 14.31 ± 14.95 m, *t* = 3.32, *p* = 0.002).

**FIGURE 3:**
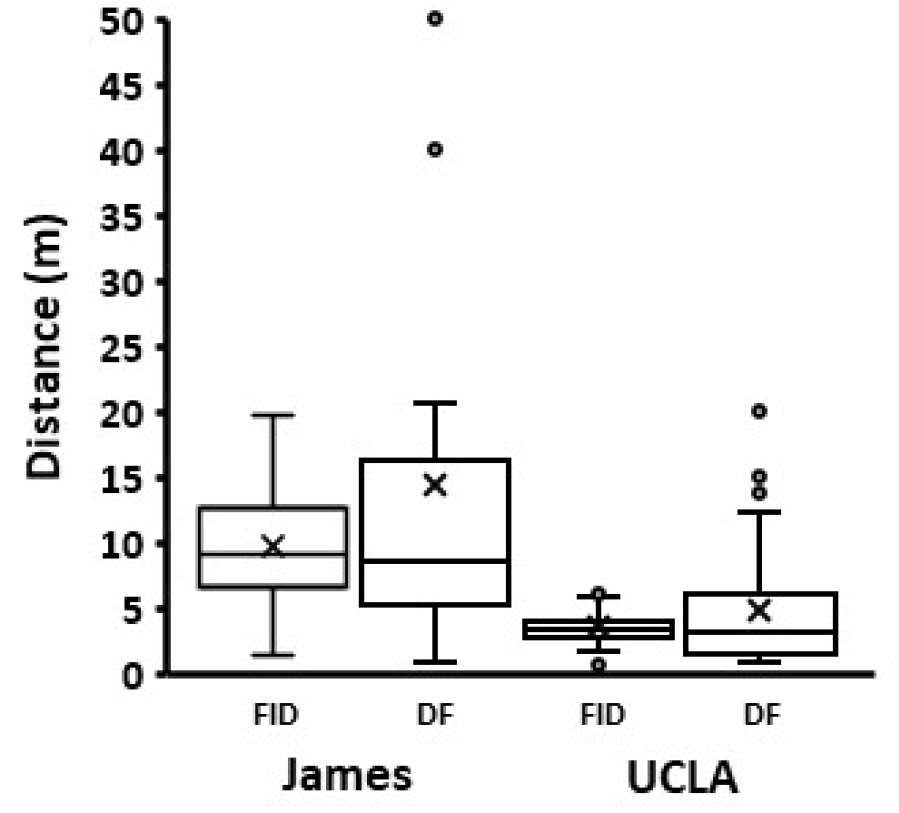
Differences in flight initiation distance and distance fled. Data are presented for the first approach a bird received at the non-urban (James, n = 22) and urban (UCLA, n = 31) sites.

Starting distances strongly influenced the first-flush FID (Fig. 4) for non-urban juncos (*F*_1,20_ = 18.84, *R*^*2*^ = 0.459, *p* < 0.001), but not significantly for urban juncos (*F*_1,29_ = 1.04, *R*^*2*^ = 0.001, *p* = 0.317). Starting distance had no significant effect on DF for either non-urban or urban birds on their first flush (non-urban: *F*_1,18_ = 0.600, *R*^*2*^ < 0.001, *p* = 0.449; urban: *F*_1,29_ = 0.063, *R*^*2*^< 0.001, *p* = 0.804).

**FIGURE 4:**
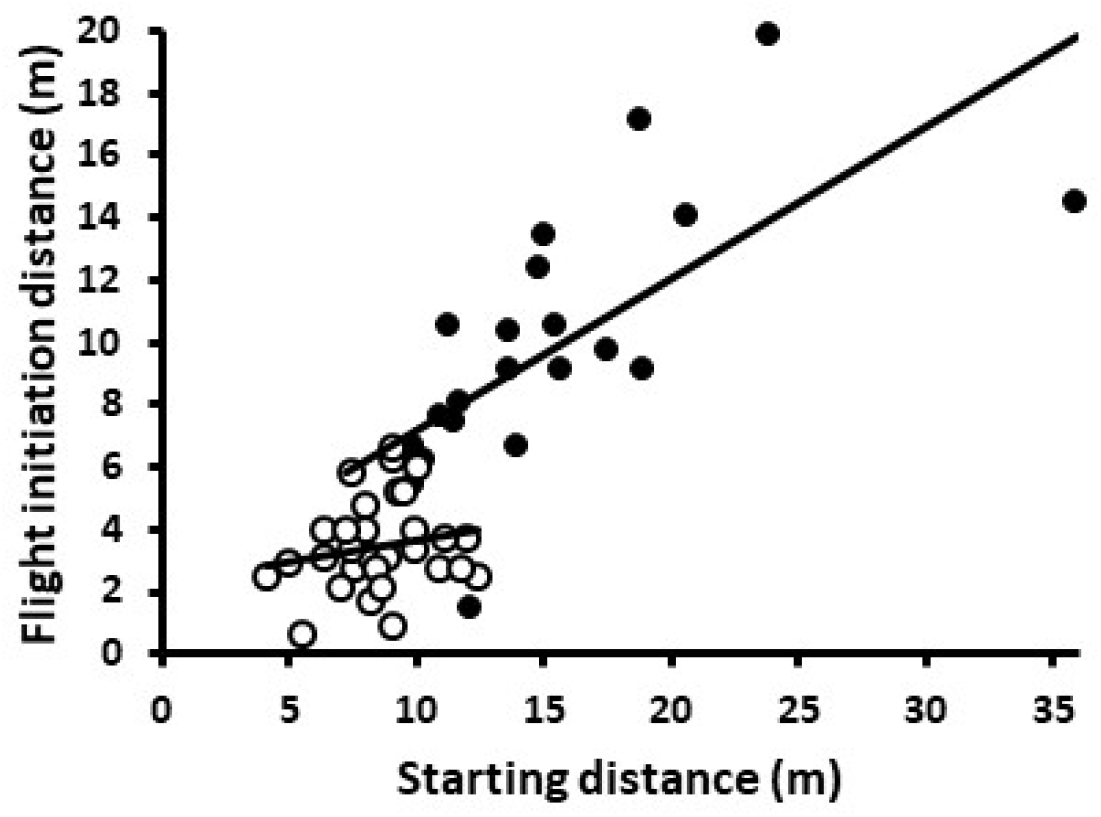
Effect of starting distance on flight initiation distance on a bird’s first flush. Open points are for urban (UCLA, n = 31) birds and solid points are for non-urban (James, n = 22) birds. Lines show the respective regressions.

## DISCUSSION

The urban juncos on the UCLA campus differed significantly from a nearby non-urban population in tolerating closer approaches by humans before moving and then moving shorter distances away. The distributions of FID at the two sites were not overlapping: of the 31 birds with the smallest mean FIDs across all approaches, 30 were from UCLA, and of the 22 birds with the largest mean FIDs, only one was from UCLA. Non-urban bird FIDs were also attuned to the starting distance at which an approach began, which suggests they place closer attention to someone coming directly towards them. In contrast, SD had no significant effect in the urban population. This pattern is consistent with urban birds evolving, under directional selection, to become more tolerant than rural birds, rather than reflecting the movement from a subpopulation of rural birds that are already intrinsically tolerant of human presence.

Another possibility is that junco behavior is sufficiently plastic that birds can adjust to the different and varying demands of urban life. Indeed, individual birds at UCLA do show behavioral plasticity in responding to repeated human approaches. However, this plasticity yields diametrically opposite outcomes. Some birds consistently habituated in FID while others appear to become sensitized as they were approached within and across days. (Because James birds could not be reliably approached multiple times within a day and then across consecutive days, it is unknown whether a similar diversity of response exists in the non-urban population.)

Furthermore, the multigenerational lag between when urban centers were first established and when juncos first colonized them, argues against such levels of behavioral plasticity being intrinsic features of junco populations everywhere (Yeh 2004; Yeh and Price 2004). Therefore, neither a specific urban-friendly genotype nor consistency in behavioral plasticity within non-urban ancestors is sufficient to account for the observed patterns in the behavior of the UCLA birds. The remaining possibility is that this and other urban junco populations are undergoing fairly rapid evolution and genetic differentiation in tolerance behavior from their non-urban ancestors (Yeh 2004; Atwell et al. 2012, 2014).

The responses of urban juncos at UCLA adds to a growing body of studies of bird behavior relative to human activity. One commonly shared characteristic across multiple species is the greater tolerance of nearby human presence in urban populations than in non-urban ones (Møller 2008, Evans et al. 2010, Samia et al. 2015, Battle et al. 2016, Cavalli et al. 2016, Vincze et al. 2016, Sprau and Dingemanse 2017). Beyond this broad pattern, however, species exhibit distinct differences in behavioral patterns. For example, as a population the UCLA juncos do not consistently habituate to being approached. This is similar to Great Tits (*Passer domesticus*: Sprau and Dingemanse 2017), but unlike House Sparrows (*Passer domesticus*: Vincze et al. 2016). However, the within-population pattern seems more similar to that reported for Burrowing Owls (*Athene cunicularia*), where studies differ as to whether or not individual birds habituate (Carrete et al. 2013, Cavalli et al. 2018). This existing across-individual variance in behavior suggests that the UCLA junco population did not arise due to differential colonization of genotypes from the ancestral non-urban population (Vincze et al. 2016), or that only certain behavioral types have segregated in the campus habitat (Sprau and Dingemnase 2017). Finally, unlike a survey across multiple bird species (Mikula 2014), we found a non-significant trend for birds being more, not less, responsive at lower pedestrian density. The suggests that the degree to which juncos view humans as a risk, they perceive it qualitatively and not quantitatively.

Another likely commonality shared with other urban populations of birds is that recent urban junco populations have likely experienced founder effects and reduced population-level genetic diversity (Møller 2010, Mueller et al. 2018). This may be creating opportunities for rapid genetic evolution (Atwell et al. 2012, 2014). The populations of Southern California juncos, therefore, appear to be an excellent system to investigate if behavioral tolerance for urban life has an evolvable genetic underpinning.

## Supporting information

Original data

## ACKNOWLEDGEMENTS

We thank Richard Hedley, Jeffrey Lee, Felisha Wong, Samuel Bressler, and Kara Lukas for assistance in the field, and Jennifer Gee, Director, and Andrea Campanella, Assistant Director of the UC James Reserve for their support. Funding was generously provided by the Santa Monica Bay Audubon Society. Eva Horna Lowell provided the Spanish language abstract.

## Author contributions statement

PJY, PN, and DTB conceived the initial idea and questions. HMS and DTB developed the methods for the experiment. HMS banded the birds and did the field work. HMS and PN analyzed the data (from programs written by DTB). HMS wrote the first draft and PN, DTB and PJY edited subsequent versions. PJY and PN contributed resources and funding to the project.

## Ethics Statement

This study was conducted under the approval of University of California - Los Angeles Animal Research Committee (protocols #2016-023-03 and 2000-147-61). All banding procedures and experimental approaches followed well-established protocols and were designed to minimize risk and disturbance to birds sampled. Junco banding was conducted under USGS Federal Bird Banding permit #23809 (to PJY) and California Department of Fish and Wildlife Scientific Collecting Permit #SCP-13549 (to HMS).

## Supporting Information

**SUPP. TABLE 1:** Information for fixed and random effects in chosen linear mixed-effects models for flight-initiation distance. Fixed-effects are mean-estimates, while random effects are variance estimates. *P*-values for fixed effects calculated using “lmerTest” (R package: “lmerTest “). LRT and *P*-values for the inclusion of random effects calculated using exactLRT (R package: “RLRsim “). Confidence intervals for fixed effects and estimates for random effects calculated using the “stats” base R package.

| Sample interval | Fixed effect | Estimate (SE) | Confidence interval (95%) | t | df | *P* |
| --- | --- | --- | --- | --- | --- | --- |
| Within-day | starting distance | 0.30 (0.10) | 0.10, 0.50 | 3.07 | 1,72 | **0.003** |
|  | Random effect | N<sub>groups</sub>, N<sub>obs</sub> | Adjusted Repeatability (SE) |  | LRT | *P* |
|  | 1\|bird | 22,72 | 0.43 (0.12) |  | 12.13 | **< 0.001** |
| Across-days | starting distance | 0.27 (0.08) | 0.10, 0.44 | 3.20 | 1,49 | **0.002** |
|  | Random effect | N<sub>groups</sub>, N<sub>obs</sub> | Adjusted Repeatability (SE) |  | LRT | *P* |
|  | 1\|bird | 23,72 | 0.43 (0.12) |  | 12.76 | **< 0.001** |

**SUPP. TABLE 2:** Information for chosen models for distance fled. For both sample durations, random effects of individual bird were found to be non-significant (via likelihood ratio test), so a model containing only fixed effects was selected. Models are generalized logistic models (GLM), with distance fled coded into near (0) and far (1), where “far” is a distance fled greater than 2 m. Confidence intervals were calculated using R software. *P*-values are produced via Wald tests.

| Sample interval | Fixed effect | Estimate<br>(logit, SE) | Estimate<br>(Odds) | Confidence interval<br>(95%, Odds) | z | *P* |
| --- | --- | --- | --- | --- | --- | --- |
| Within-day | FID | 0.44 (0.19) | 1.55 | 1.09, 2.33 | 2.30 | **0.02** |
| Across-days | FID | -0.86 (0.26) | 0.42 | 0.23, 0.67 | -3.25 | **0.001** |
|  | distance to cover | -0.30 (0.17) | 0.74 | 0.52, 1.0 | -1.77 | 0.08 |

**SUPP. TABLE 3:** Comparison of models incorporating samples collected within the first sample day for each urban individual. Fixed effects were chosen based on AIC. Sample iteration (Trial within day) does not appear to significantly improve model fit. Model m0.1 was selected as the best model for this subset of the data.

| Model Name | ΔAIC | Model Formula | Comparison | *P* |
| --- | --- | --- | --- | --- |
| m0 | 6.5 | fid ~ (1\|bird) | NA | NA |
| **m0.1<sup>a</sup>** | **0.0** | **fid ~ starting distance + (1\|bird)** | **m0.1,m0** | **0.003** |
| m1 | 0.2 | fid ~ starting distance + trial + (1\|bird) | m1,m0.1 | 0.19 |
| m2 | 2.7 | fid ~ starting distance + trial + (1 + trial\|bird) | m2, m0.1 | 0.35 |
| m2 | see above | see above | m2,m1 | 0.46 |

<sup>a</sup>The AIC value for the selected model for FID within-day is 274.9.

**SUPP. TABLE 4:** Comparison of models constructed from samples over long time scales, incorporating first flushes across sample days. Fixed effects were chosen based on AIC. Sample iteration (Day) was not found to have a significant influence on the quality of the model fit. Ultimately, model m0.1 was chosen based on AIC and comparison with other models using likelihood ratio tests.

| Model Name | ΔAIC | Model Formula | Comparison | *P* |
| --- | --- | --- | --- | --- |
| m0 | 7.8 | fid ~ (1\|bird) | NA | NA |
| **m0.1<sup>b</sup>** | **0.0** | **fid ~ starting distance + (1\|bird)** | **m0.1, m0** | **0.002** |
| m1 | 2.0 | fid ~ starting distance + day + (1\|bird) | m1, m0.1 | 0.81 |
| m2 | 5.9 | fid ~ starting distance + day + (1 + day\|bird) | m2, m1 | 0.95 |

<sup>b</sup>The AIC value for the selected model for FID across-days is 267.

**SUPP. TABLE 5:** Comparison of models constructed to examine distance fled over short time scales. We selected a model lacking random effects of individual birds based on likelihood ratio tests and AIC. Distance fled was binned into two levels, near (0) and far (1), where “far” is a distance fled greater than 2 m. Fixed effects were chosen adding and removing predictor variables until the lowest AIC was reached.

| Model Name | ΔAIC | Model Formula | Comparison | *P* |
| --- | --- | --- | --- | --- |
| m0 | 6.11 | nearfar ~ (1\|bird) | N/A | N/A |
| m0.1 | 2 | nearfar ~ fid + (1\|bird) | m0.1,m0 | 0.01 |
| m1 | 1.94 | nearfar ~ fid + trial + (1\|bird) | m1,m0.1 | 0.15 |
| m2 | 5.94 | nearfar ~ fid + trial + (1 + trial\|bird) | m2, m0.1 | 0.56 |
| m2 | see above | see above | m2,m1 | 1 |
| **m0.1glm<sup>c</sup>** | **0.00** | **nearfar ~ fid** | **m0.1,m0.1glm** | **1** |

<sup>c</sup>The AIC value for the selected model of DF within-day is 71.34.

**SUPP. TABLE 6:** Model comparison for DF from samples collected across days. As with within-day data, we selected a model with no individual effect based on a combination of likelihood ratio tests and AIC. Fixed effects were selected by adding and removing predictor variables and comparing AIC. While distance to cover is not clearly significant (see Wald test result in Supplementary Table 2), likelihood ratio tests suggest that its inclusion in both mixed-logistic and logistic models improve the fit of the models (likelihood ratio test, *P* = 0.02; logistic Δresidual deviance = −3.87, *P* = 0.05).

| Model Name | ΔAIC | Model Formula | Comparison | $P$ |
| --- | --- | --- | --- | --- |
| m0 | 15.93 | nearfar ~ (1\|bird) | NA | NA |
| m0.1 | 0.13 | nearfar ~ distance to cover + fid + (1\|bird) | m0.1,m0 | < 0.001 |
| m1 | 1.91 | nearfar ~ distance to cover + fid + day + (1\|bird) | m1,m0.1 | 0.64 |
| m2 | 4.58 | nearfar ~ distance to cover + fid + day + (1 + day\|bird) | m2,m0.1 | 0.67 |
| m2 | see above | see above | m2,m1 | 0.51 |
| **m0.1glm<sup>d</sup>** | **0.00** | **nearfar ~ distance to cover + fid** | **m0.1, m0.1glm** | **0.17** |

<sup>d</sup>The AIC value for the selected model of DF across-days is 67.92.

